# Reductions in Retrieval Competition Predict the Benefit of Repeated Testing

**DOI:** 10.1101/292557

**Authors:** Nicole S. Rafidi, Justin C. Hulbert, Paula Pacheco, Kenneth A. Norman

## Abstract

Repeated testing leads to improved long-term memory retention compared to repeated study, but the mechanism underlying this improvement remains controversial. In this work, we test the hypothesis that retrieval practice benefits subsequent recall by reducing competition from related memories. This hypothesis implies that the degree of reduction in competition between retrieval practice attempts should predict subsequent memory for practiced items. To test this prediction, we collected electroencephalography (EEG) data across two sessions. In the first session, participants practiced selectively retrieving exemplars from superordinate semantic categories (high competition), as well as retrieving the names of the superordinate categories from exemplars (low competition). In the second session, participants repeatedly studied and were tested on Swahili-English vocabulary. One week after session two, participants were again tested on the vocabulary. We trained a within-subject classifier on the data from session one to distinguish high and low competition states.We then used this classifier to measure competition across multiple retrieval practice attempts in the second session. The degree to which competition decreased for a given vocabulary word predicted whether that item was subsequently remembered in the third session. These results are consistent with the hypothesis that repeated testing improves retention by reducing competition.

## Introduction

How can learners maximize their retention of important information while minimizing time spent studying? Compared to repeatedly testing oneself, passively reviewing the to-be-learned material may seem like a relatively painless and effective way to confirm what is known and strengthen what is not. However, numerous studies have found that repeated testing leads to better long-term retention than repeated study^1,2^. In one notable experiment, Karpicke and Roediger^3^ trained participants to the point where they could recall items correctly, and then manipulated (as part of the same learning session) whether participants received additional memory tests or study trials for those items. When Karpicke and Roediger tested memory one week later, they found that delayed recall was much better in the additional-testing condition than the additional-study condition (80% recall after testing vs. 36% recall after restudy).

How can we explain this huge benefit of testing on delayed retention? A number of accounts for the testing effect have been offered in the literature, with varying degrees of support from neural and behavioral data^1,2^. One theory posits that the additional mental effort required for selective retrieval may reflect a desirable difficulty that enhances retention^4,5^. The close match between the cognitive processes involved during practice tests and the final test (i.e., transfer-appropriate processing) may also contribute to the benefit^6^. Another possibility is that searching for the correct response on a test leads to the elaboration of semantic associations through which the target material can later be accessed^7^. Yet another potential explanation is that repeated testing serves to reduce competition from related memories, thereby improving access to the target memory^8^.

Speaking to this last possibility, a large body of evidence highlights how selective retrieval(but not passive restudy) impairs the accessibility of competing memories. For example, repeatedly retrieving “SCOTCH” when cued with “ALCOHOL-S” impairs later recall of other alcoholic drinks (like “VODKA”) relative to unpracticed baseline categories. Simply reviewing “ALCOHOL-SCOTCH” facilitates recall of “SCOTCH,” but it is not associated with a corresponding impairment to the accessibility of “VODKA.” Since the first report of retrieval-induced forgetting (RIF)^9^, a number of explanations for this phenomenon have been posed^10–12^. Most commonly it has been attributed to the inhibition of competing memories^9, 13–16^. Indeed, the results of a recent meta-analysis largely support the inhibitory account^17^, which notably has been implemented in a neural network model^18^ that has helped to explain and predict various neural and behavioral results 1^9^.

Should weakening of competitors be responsible for RIF, one might expect overall levels of competition to decrease over successive retrieval attempts; the relative reduction in competition should, in turn, predict competitor forgetting on a later test. Functional magnetic resonance imaging (fMRI) data support this prediction: One study found that, across multiple retrieval practices, decreased activity in anterior cingulate cortex (a region thought to reflect conflict between possible responses) predicted the degree to which participants showed an RIF effect; these particular changes in brain activity were selective to successful retrieval practice^20^. Another study found that, as unwanted intrusions from competing memories declined over the course of retrieval practice, so too did the extent to which their corresponding neural patterns were reactivated – all in a manner that predicted competitor forgetting^21^.

While the aforementioned results show that testing reduces competition, it has not yet been shown that this competition reduction supports the beneficial effects of testing on long-term retention of the tested items^2^. Here, we sought to test this hypothesis by obtaining a neural measure of competition, tracking its decline across retrieval practice attempts, and then relating this decline to subsequent memory for the practiced items (one week later).

To neurally measure competition, we leveraged prior work by Hanslmayr and colleagues that showed strong EEG differences for “high competition” retrieval(in which participants selectively retrieved one of many exemplars from a superordinate semantic category) as compared to “low competition” retrieval (in which participants retrieved the superordinate category given an exemplar)^22^. In our study, we trained a within-subject EEG pattern classifier on these two conditions to distinguish between high and low states of competition. This part of the experiment (Session 1) can be viewed as a kind of “competition localizer” that uncovers the neural signature of competition in EEG data.

After this initial “competition localizer” phase, we used a variant of the paradigm initially used by Karpicke and Roediger to show benefits of testing^3^: Participants were shown a set of Swahili-English word pairs and then were given multiple cycles of study and retrieval practice to learn these word pairs. In the original Karpicke and Roediger experiment, they compared several conditions that varied the amount of restudy and retrieval practice that occurred; in our study, our primary interest was in variance in competition *within* the retrieval practice condition, so we did not manipulate learning conditions across participants – we only included a single condition where participants received a fixed amount of retrieval practice and restudy, regardless of whether an item had been recalled successfully.We refer to this part of our experiment (Session 2) as the Swahili Learning Task.

We recorded EEG during retrieval practice trials in Session 2.We applied the competition classifier (trained on Session 1) to these trials, thereby obtaining a trial-by-trial measure of competition. Lastly, we had participants return one week after Session 2 for another session (Session 3) where they were cued with Swahili words and asked to recall the associated English word. A schematic of the experiment with all three sessions can be found in Fig. 1.

**Figure 1.**
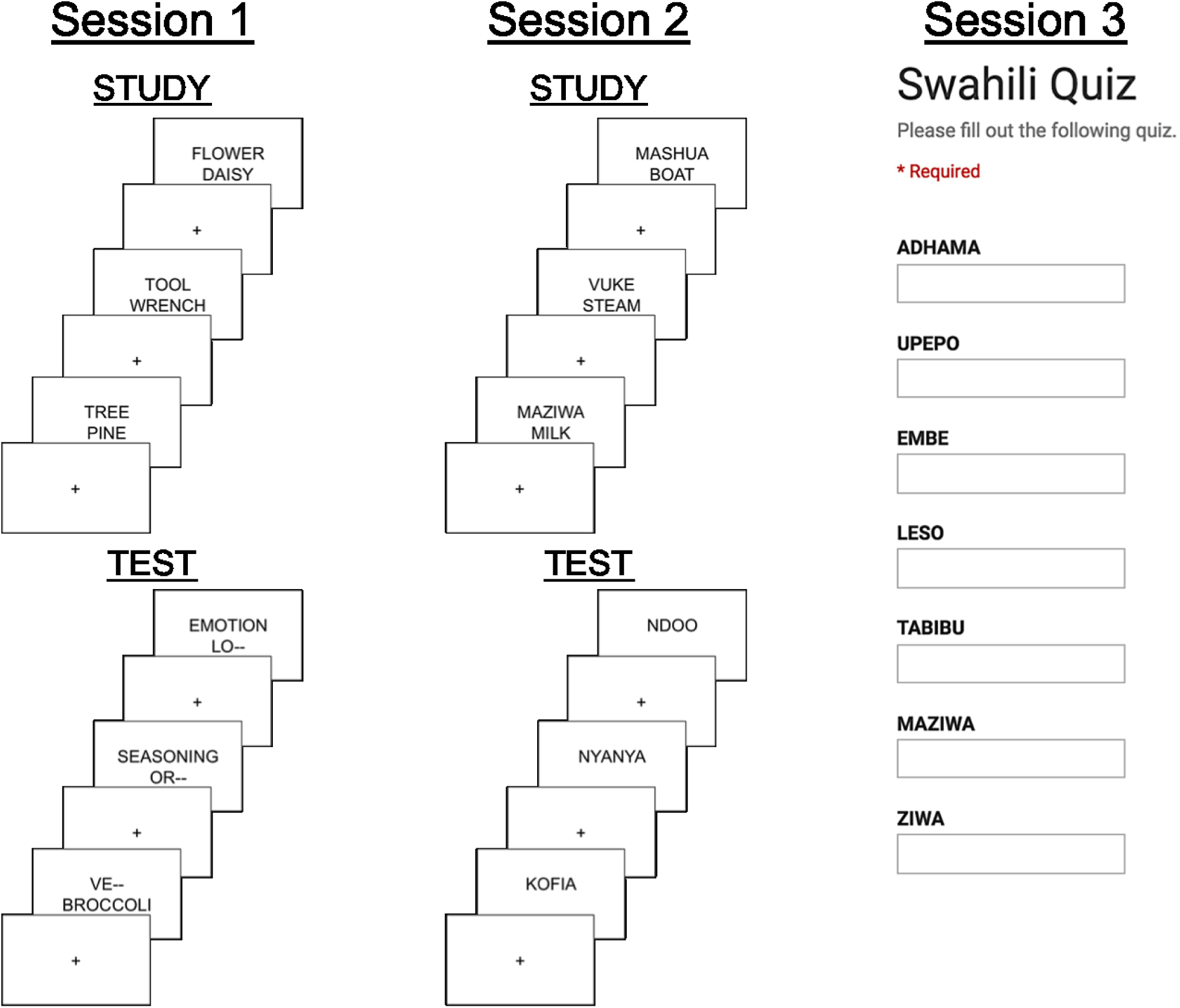
Schematic of experimental design. Session 1 consisted of a single study block of English category-exemplar pairs, followed by four test blocks contrasting high- and low-competition cued recall. In high-competition trials, the category was provided along with the first two letters of the exemplar (e.g. “TOOL-WR–”), and in low-competition trials, the exemplar was provided along with the first two letters of the category (e.g. “TO– WRENCH”). Subjects were asked to silently finish the partially blanked-out word (the blank was not diagnostic of the length of the word). Session 2 consisted of alternating study and test blocks (four total of each) for Swahili-English word pairs. During test blocks, subjects silently thought of their response and then typed it in after a 2 second delay. The questions in the test round did not progress unless the participant hit “enter”. EEG data were collected during Sessions 1 and 2. Session 3 was a simple cued recall quiz on the items studied in Session 2, with the Swahili words presented as cues (no EEG was recorded during Session 3).

For each word pair in Session 2, we computed the difference in the measured level of competition between the first successful retrieval practice trial for a given item and the last retrieval practice trial for that item; we call this measure “competition drop”. We chose the first *successful* retrieval practice trial for this measure because – like Karpicke and Roediger ^3^ – we were interested in the learning processes that take place *after the first successful recall attempt* that determine whether or not the memory will be retained one week later. We hypothesized that the size of the competition drop value (i.e., the degree to which competition was reduced for this pair) would predict whether participants successfully recall that pair in Session 3. Our results support this hypothesis, as described below.

## Results

### Behavioral Results

Average performance across participants is shown in Fig 2a. On average, performance improved over each of the four retrieval practice trials (testing rounds) in Session 2 (R1-R4) but dropped between the last Session 2 testing round (R4) and the Session 3 test (T). The average performance on Session 3 was near 50%, which gives an (approximately) balanced number of remembered and forgotten items for analyzing which factors (in Session 2) predict subsequent recall in Session 3. Performance in Session 3 varied over subjects, as shown by the histogram of subject performance values in Fig 2b. While the majority of subjects retained at least 50% of the Swahili-English word pairs, a substantial number retained less than that.

**Figure 2.**
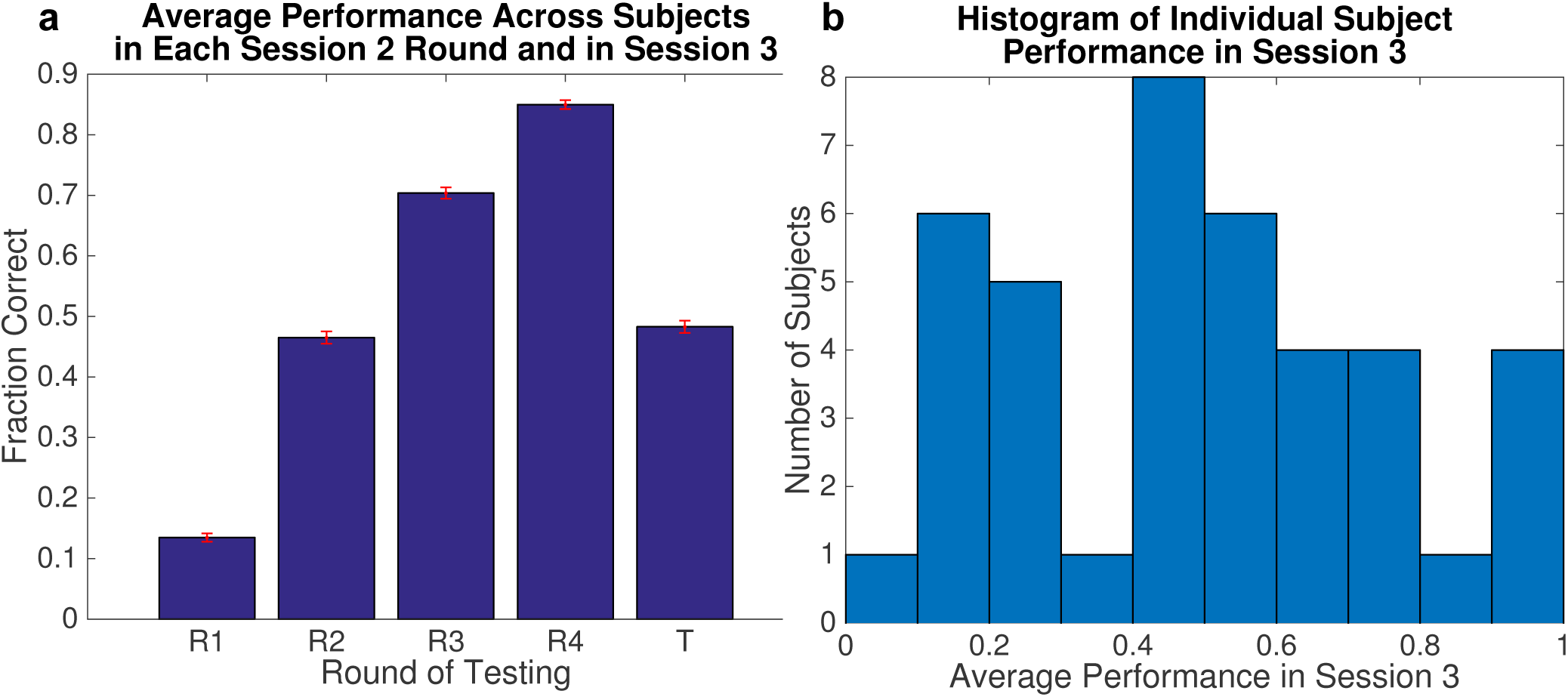
Behavioral performance summary. a. Behavioral performance on testing rounds in Session 2 (R1-R4) and Session 3 (T) Each bar was computed by taking the mean over the pool of all trials from all subjects. The error bars show standard error. On average, performance improved over each round in Session 2, and approximately half the words were remembered in Session 3. **b. Histogram of individual subject recall performance in Session 3.** Most subjects were able to retain some of the words from Session 2, and the majority remembered at least 50%, but there was a substantial group of low-performing subjects who remembered less than 50% of the studied words.

We can use the behavioral data from Session 2 to distinguish between subsequently remembered items (i.e., items correctly recalled in Session 3) and subsequently forgotten items. Fig 3 shows the average recall performance on each Session 2 testing round, grouped by performance on the Session 3 test. The blue and yellow bars show the average performance on subsequently forgotten and remembered items, respectively. Unsurprisingly, items that were better remembered in Session 3 were also better remembered in Session 2.

**Figure 3.**
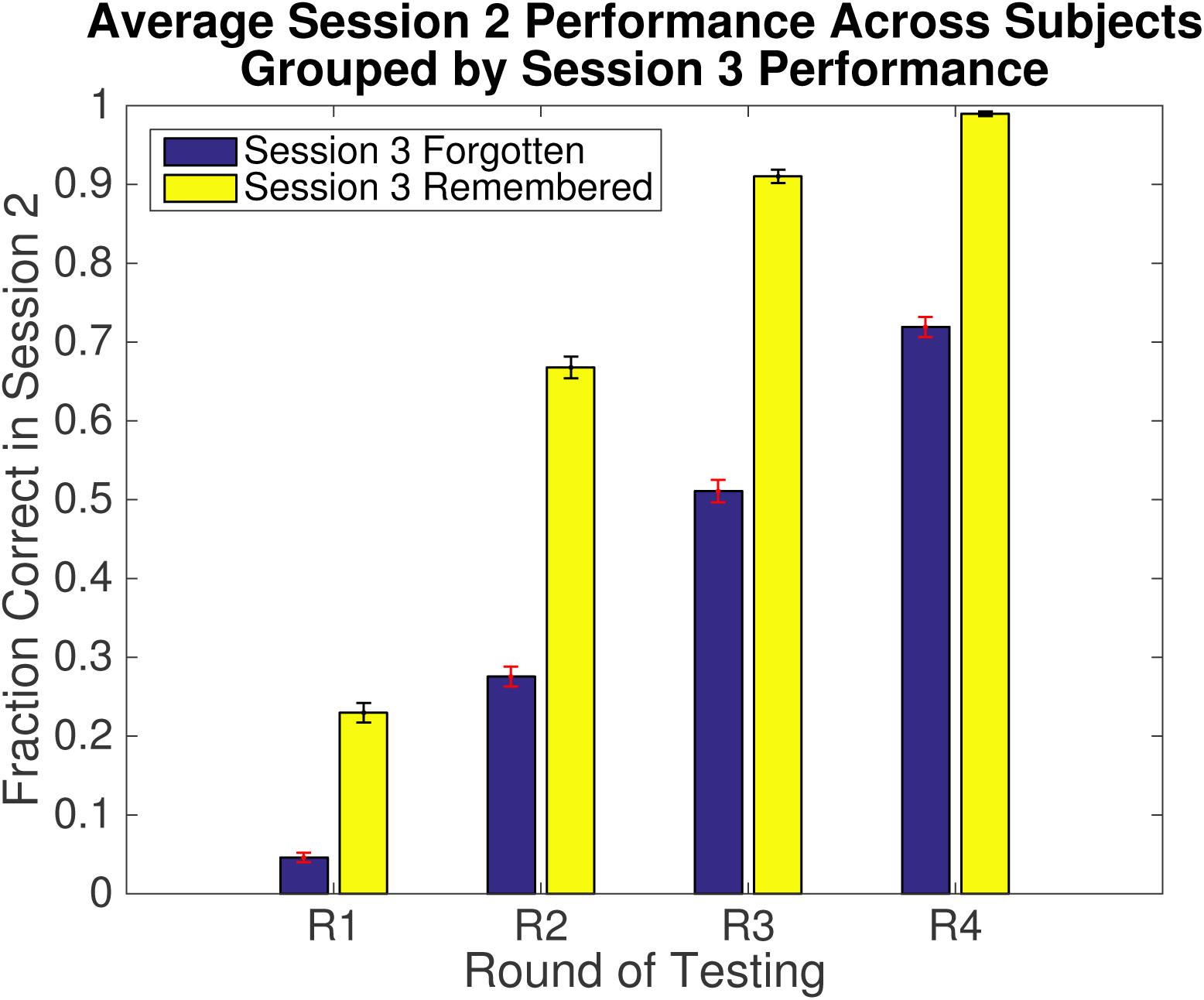
Behavioral performance on Session 2 testing rounds (R1-R4), grouped by Session 3 performance. Performance on each round was divided into two averages: items that were subsequently forgotten in Session 3 (blue) and items that were subsequently remembered (yellow). As in Fig 2, each bar indicates the mean performance over all subjects and items, and error bars show standard error.

Next we examined how particular *patterns* of correct vs. incorrect recall (across trials) in Session 2 were related to recall in Session 3.For the purpose of this analysis, we encoded the Session 2 responses as 4-dimensional binary vectors, where a 1 in the *i^th^* position indicates that the participant responded correctly in the *i^th^* round of Session 2. As an example, a word for which the participant responded correctly in all but the first round would be encoded as 0111.

The result is shown in Fig 4. Some response configurations were much more likely than others; in general, once a participant was able to recall an item correctly, they continued to be able to do so in subsequent rounds. The more rounds in which the participant answered correctly, the more likely the participant was to remember that item in Session 3. Using a logistic regression classifier on the trials pooled over all subjects, we achieved 64% classification accuracy using recall accuracy for the four test rounds (in Session 2) as input features and subsequent recall accuracy (in Session 3) as the dependent measure. Chance is 50%. A permutation test with 1000 permutations yielded *p* < 0.001, demonstrating that Session 2 behavior was very significantly predictive of Session 3 recall.

**Figure 4.**
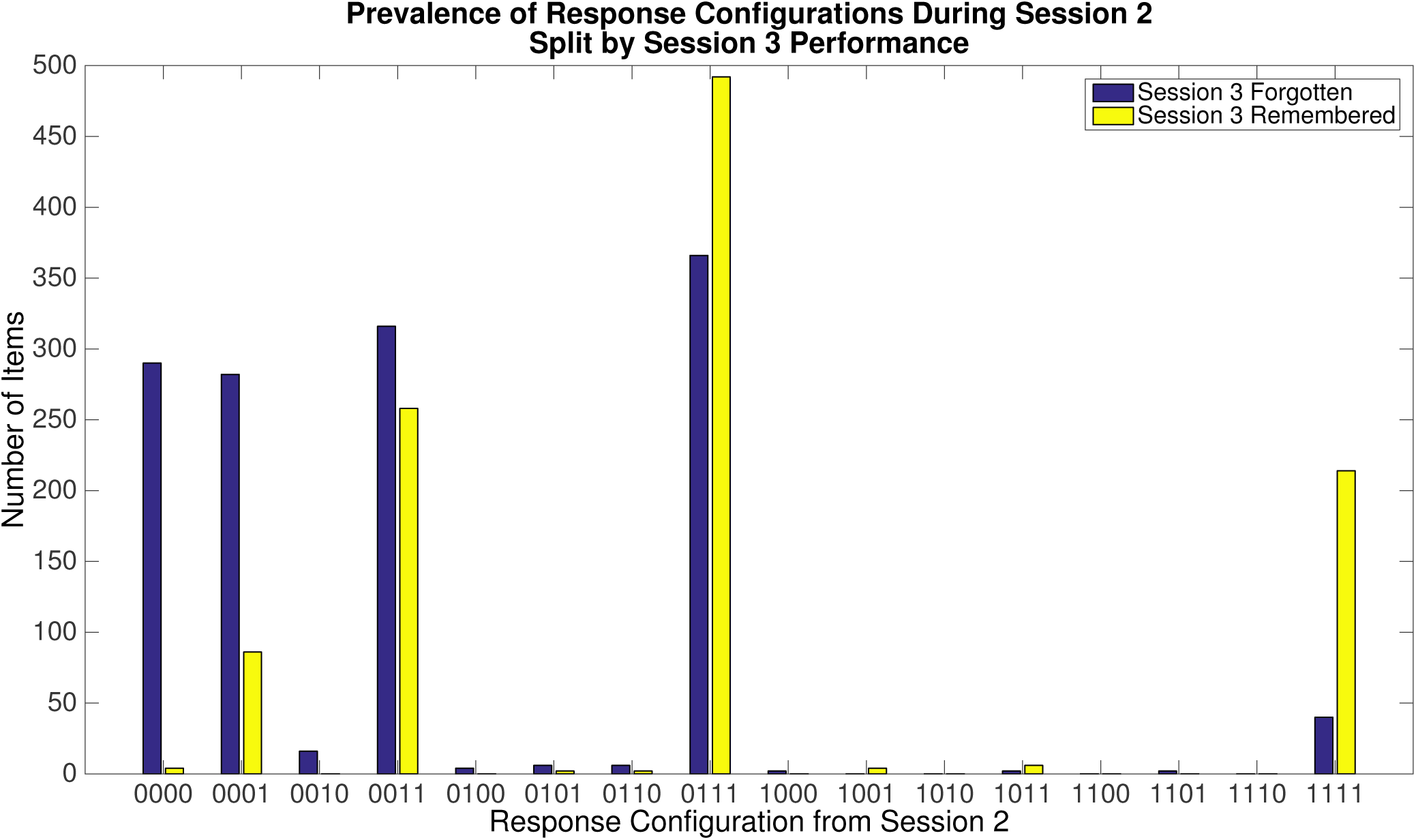
Predictive power of Session 2 behavioral data. Each pair of bars shows the number of items (pooled over subjects) with the labeled response configuration, where a 1 in the *i^th^* position indicates that the subject responded correctly in the *i^th^* testing round of Session 2. Blue bars show items that were subsequently forgotten, while yellow show subsequently remembered items in Session 3.

## EEG Analysis

### Classifying High and Low Competition Neural States

We trained a classifier to distinguish between high and low competition trials from the Session 1 EEG data. Prior work using a similar paradigm found EEG signals that discriminated between high and low competition trials during the first 500 ms post-stimulus-onset^22^; however, we did not have a strong *a priori* hypothesis about exactly when (within this window) competitive states would be most distinguishable, and we did not want to rule out the possibility that slightly later time points would also discriminate. As such, we applied the classifier in a sliding window fashion, retraining and testing for each time point independently. The features given to the classifier were the voltages at each of the 64 EEG channels, averaged over a 50ms window. We were able to successfully decode over a large period of time post-stimulus-onset, shown in Fig 5. Significantly-above-chance decoding started at the 60ms window (encompassing data from 60ms to 110ms post-stimulus-onset), and tapered off at 640ms post stimulus onset. Peak competition decoding accuracy occurred at 220ms post-stimulus-onset. For subsequent analysis, we chose the segment of time for which decoding was significant and for which accuracy was greater than one standard deviation (SD) above chance. This encompasses the time window from 80ms-to-600ms post-stimulus-onset (note that accuracy dips below 1-SD-above-chance for a brief period between 100 to 200ms; we decided to keep these timepoints in the analysis rather than introducing a discontinuity in the time window of interest).

**Figure 5.**
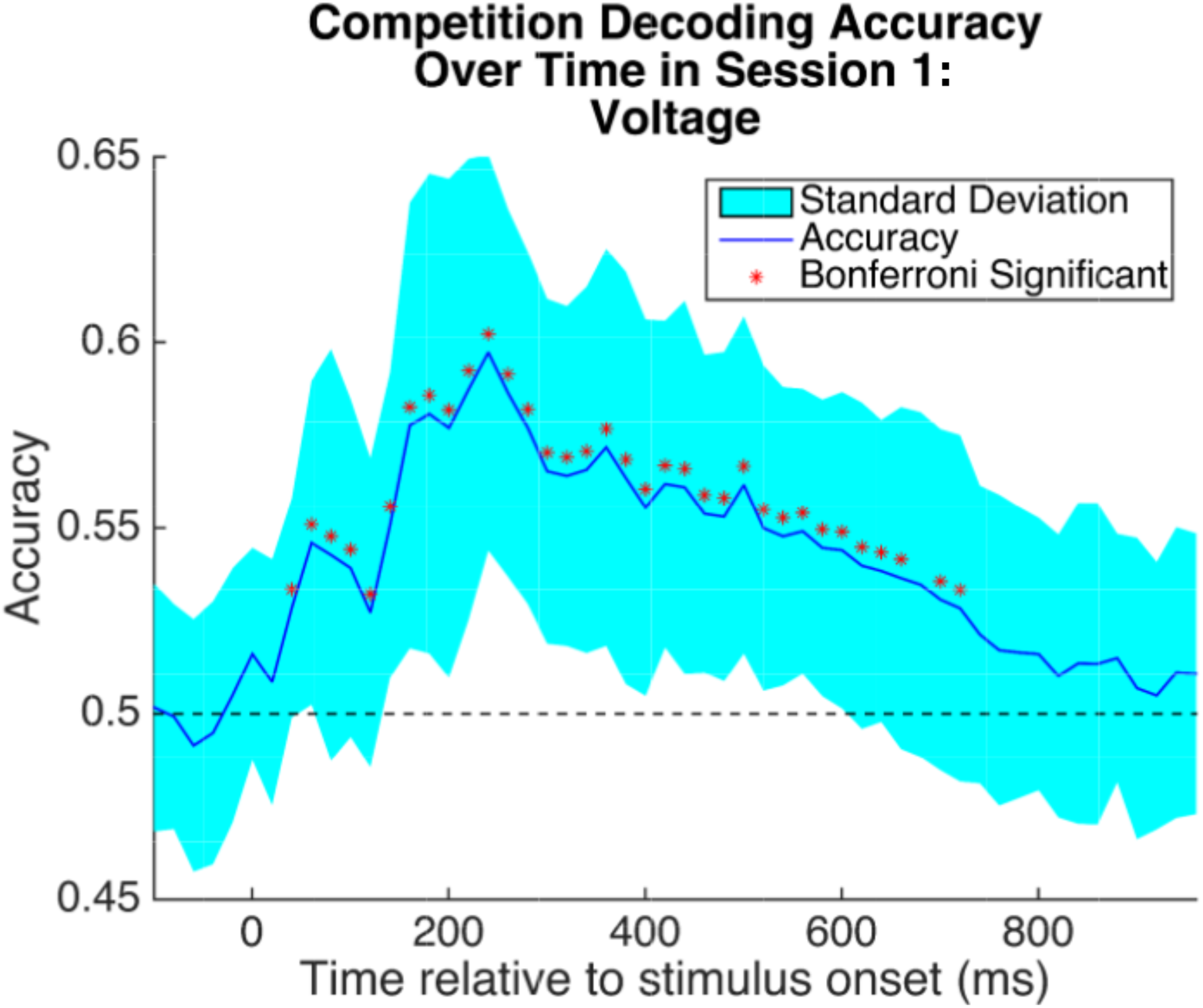
Competition decoding accuracy over time in session 1. 5-fold cross-validation accuracy at each timepoint for distinguishing high- and low-competition retrieval states, averaged across subjects. Each point used the average voltage over a 50ms window (starting at that time point) at each channel as features for classification. Points are spaced 20ms apart. The shaded region gives the standard deviation across the subject means. Chance is 50%. Stars indicate time points that were significant after per-timepoint permutation test and Bonferroni correction at 0.001 significance threshold.

To evaluate whether the neural representation of competition remained constant throughout the significantly decodable period, we computed a representational dissimilarity matrix (RDM)^23^ using the learned weight vectors at each time point. The result is shown in Fig 6. Over time, there seemed to be two distinct neural representations that were predictive: one early, from 100-200ms post-stimulus-onset, and one late, from 300-600ms post-stimulus-onset. We examined the difference between these representations by transforming the weight vectors into pattern maps using the technique described Haufe et. al^24^.We then averaged these maps within each of the two time periods of interest and made topographic plots. The results are shown in Supplementary Figure S1.

**Figure 6.**
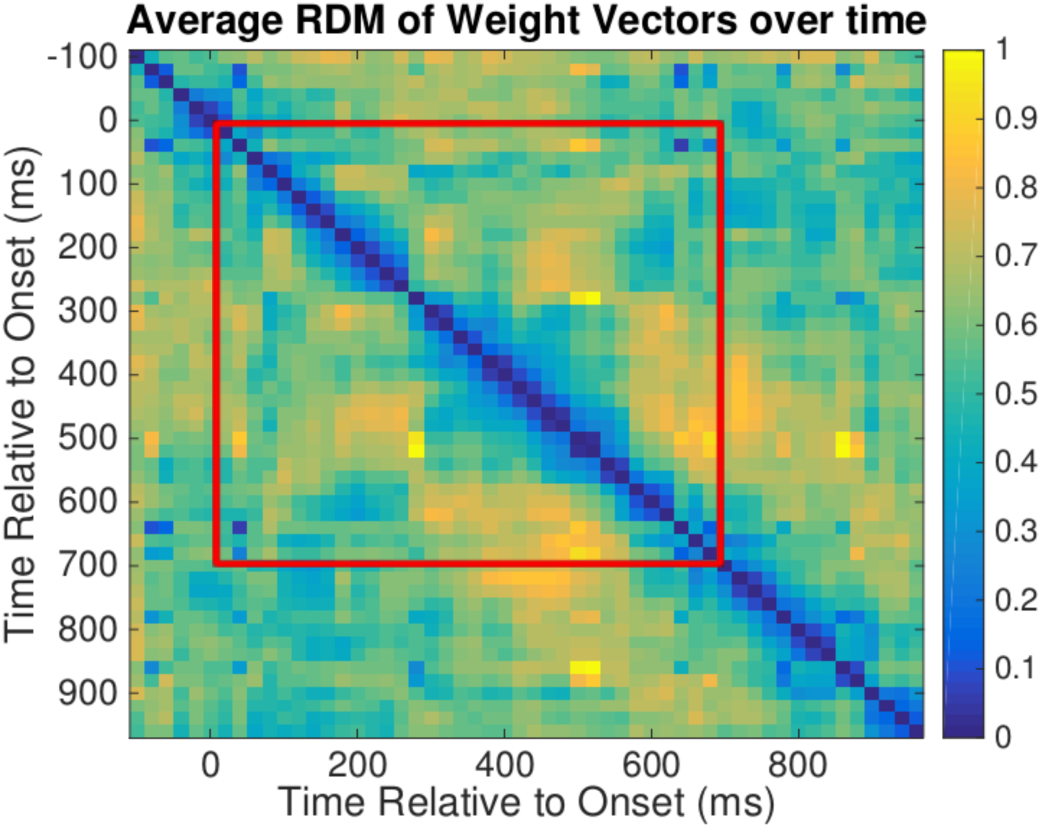
Representational dissimilarity matrix (RDM) of competition weight vectors. RDM using cosine distance to distinguish between learned logistic regression weight vectors at each time point in the competition localizer task. The portion captured in red corresponds to significant decoding window from Fig 5. Blue indicates that two time points are more similar, whereas yellow indicates that they are different.

**Figure S1.**
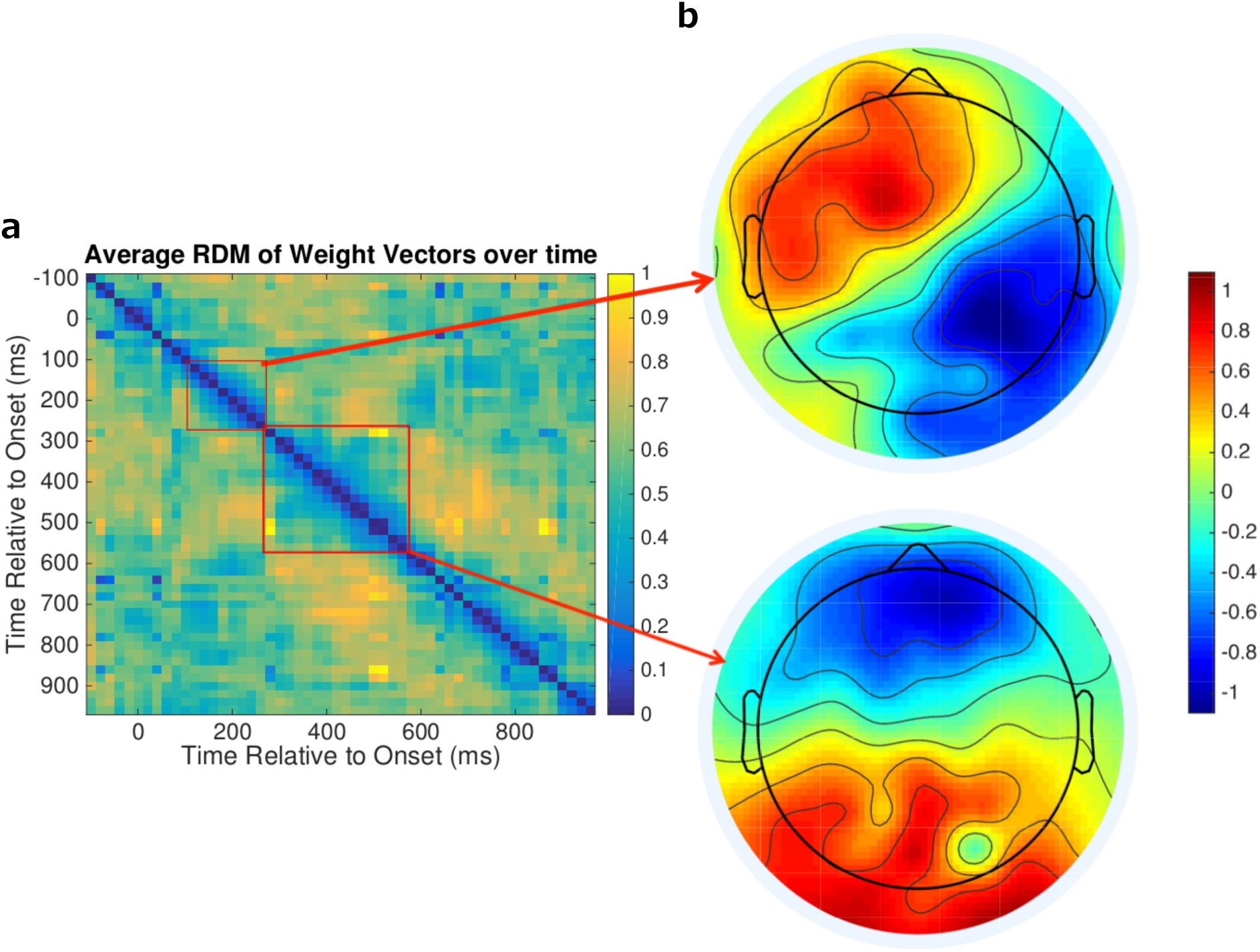
a. RDM of competition weight vectors Taken from Fig. 6b. **Topographic pattern maps** Weight matrices from the red captured regions in panel **a** were transformed into importance maps^24^ and then averaged to create the two topographic plots. Pattern values further from 0 indicate a stronger relationship between that feature and the classification task.

### Predicting Subsequent Memory from the Swahili Learning Task

For each competition-localizer time point from 80ms to 600ms post stimulus onset, we re-trained the classifier on all of the competition data from Session 1 and applied it to the EEG data from Session 2, again within-subject and in a sliding window fashion, generating a grid of competition values for each retrieval practice round (where the grid is defined by which time points post-stimulus-onset were used for training vs. testing). For each item, we computed a measure of *competition drop* by subtracting the item’s competition value for the last round (R4) from the competition value for the round for which the participant first answered correctly.We then used this measure of competition drop to attempt to distinguish between items that were subsequently remembered vs. forgotten in Session 3 (this analysis was run separately for each grid point). The initial comparison is shown in Fig 7.We had no *a priori* hypothesis about when in time our effect would occur, so we computed this grid search and then corrected for multiple comparisons over the grid.

**Figure 7.**
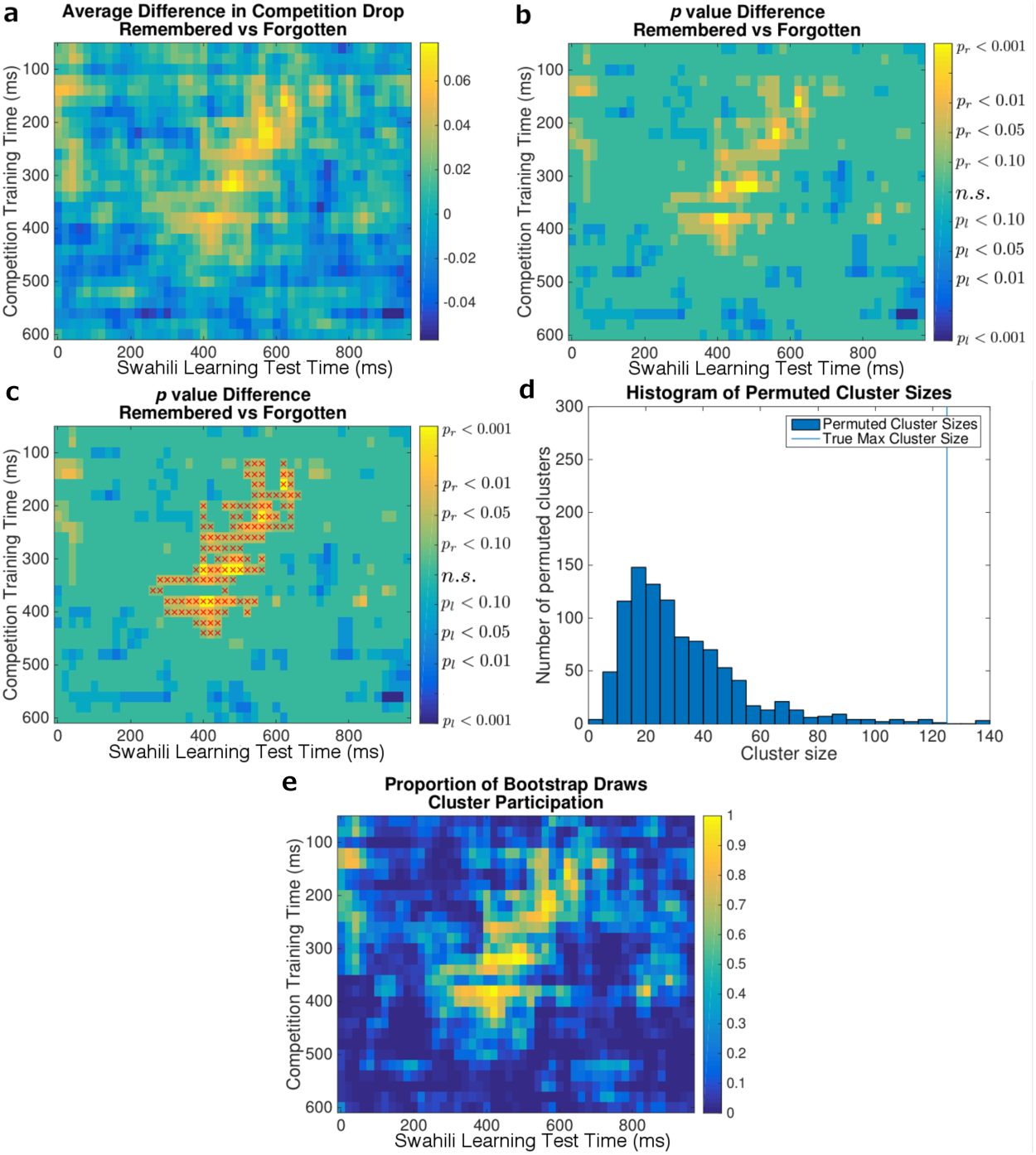
Competition drop predicts subsequent memory. We tested classifiers trained on high-accuracy timepoints from Session 1 on all post-stimulus-onset time points during the Swahili Learning Task in Session 2. This crossing of training and testing time points defines a grid. For each item and each grid square, we computed the *competition drop*: the decrease in competition across successive retrieval trials in the Swahili Learning Task (see text for details). We hypothesized that larger competition drop values in Session 2 would predict better recall in Session 3; we tested all grid squares and corrected for multiple comparisons using a cluster permutation procedure. **a. Average difference in competition drop between remembered and forgotten items.** Yellow indicates that remembered items exhibited a larger competition drop than forgotten items. Blue indicates the opposite. **b. One-tailed t-test result of the difference in competition drop between remembered and forgotten items.** Grid squares for which the drop was larger for remembered items, i.e., where a right-tailed test (*p_r_*) achieved significance at *p_r_ <* 0.10, are yellow; grid squares for which the drop was larger for forgotten items, i.e., where a left-tailed test (*p_l_*) achieved significance at *p_l_* < 0.10, are blue. The right-tailed test corresponds to our hypothesis that a drop in competition is predictive of subsequent memory. **c. Clusters that were significantly predictive of subsequent memory.** Same base plot as in **b** with the surviving cluster from permutation testing labeled with red circles. This cluster was significant at *p* = 0.003. **d. Cluster permutation test histogram.** The vertical line indicates the surviving cluster size achieved using the true labels. Each draw in the histogram represents largest cluster observed in a given permutation of the data. **e. Subject bootstrap results.** Subjects were sampled with replacement, and on each draw the cluster analysis was re-run. The color scale on this plot represents the proportion of bootstrap draws in which a point was included in a significant cluster. A high frequency (yellow) indicates the the point was highly consistent across subjects.

Figure 7 a and b show the uncorrected result of predicting subsequent memory using the competition drop metric. Results that are consistent with our hypothesis that a larger competition drop predicts successful learning are shown in yellow. Results consistent with the opposite effect are shown in blue. Note the large swath of timepoints consistent with our hypothesis. In general, classifiers trained on early Session 1 time points (i.e., the top half of the grid) were most predictive of subsequent memory performance.

We determined the (multiple-comparisons-corrected) significance level of these results using a 2-stage process. We first computed an (uncorrected) *p* value for each grid square by comparing the competition drop scores for subsequently-remembered vs. subsequently-forgotten items using a *t*-test (for this test, we pooled all items across all subjects). Next, we thresholded the *p* values from this t-test at *p* < 0.10. To correct for multiple comparisons, we used a cluster permutation test. For this test, we computed the size of the largest cluster of contiguous grid squares where *p* < 0.10; we then compared this to a null distribution of maximum cluster sizes that we generated by permuting the assignment of items to the subsequently-remembered vs. forgotten classes; this analysis yields a corrected family-wise error *p* value. The large central cluster was the only cluster to achieve a family-wise *p* < 0.05, (*p* = 0.003), marked with red dots in Fig 7c. The histogram of cluster sizes is shown in Fig 7d.

To establish the reliability of this cluster over the participant population, we computed a participant-level bootstrap (i.e.,resampling participants with replacement) and recorded the proportion of bootstrap draws for which a given grid square was included in a significant cluster. The points in the large cluster were present in nearly all bootstrap draws, as shown in Fig 7e.

These results show that the competition drop measure was strongly predictive of subsequent memory. Specifically, items for which there was a large drop in competition level from the first correct round to the last round were much more likely to be subsequently remembered. However, it is unclear from the result shown in Fig. 7 whether the competition drop metric computed from the Session 2EEG data provides additional predictive power over the Session 2 behavioral recall data alone. To address this, we re-ran the analysis on subsets of items where Session 2 behavioral recall performance is held constant. Based on Fig. 4, we selected items with 0011 and 0111 response configurations because these subsets both contained a large number of items and yielded a relatively even split between subsequently remembered and forgotten items. The result is shown in Supplementary Figure S2. The results split by behavior (panels b and c) are qualitatively similar to both the original result (panel a) and to each other. Family-wise significance was more difficult to achieve for these sub-analyses, given the drastic reduction in items (574 for 0011 and 858 for 0111, as compared with 2170 items total); the resulting family-wise *p* values were*p* = 0.01 for 0011 items and *p* = 0.17 for 0111 items. Having said this, it is striking how similar the results were for the 0011 and 0111 analyses, given that they were computed on two entirely distinct subsets of the data — these results can thus be considered to be an internal replication of our main result, modulo the power issues noted above.

**Figure S2.**
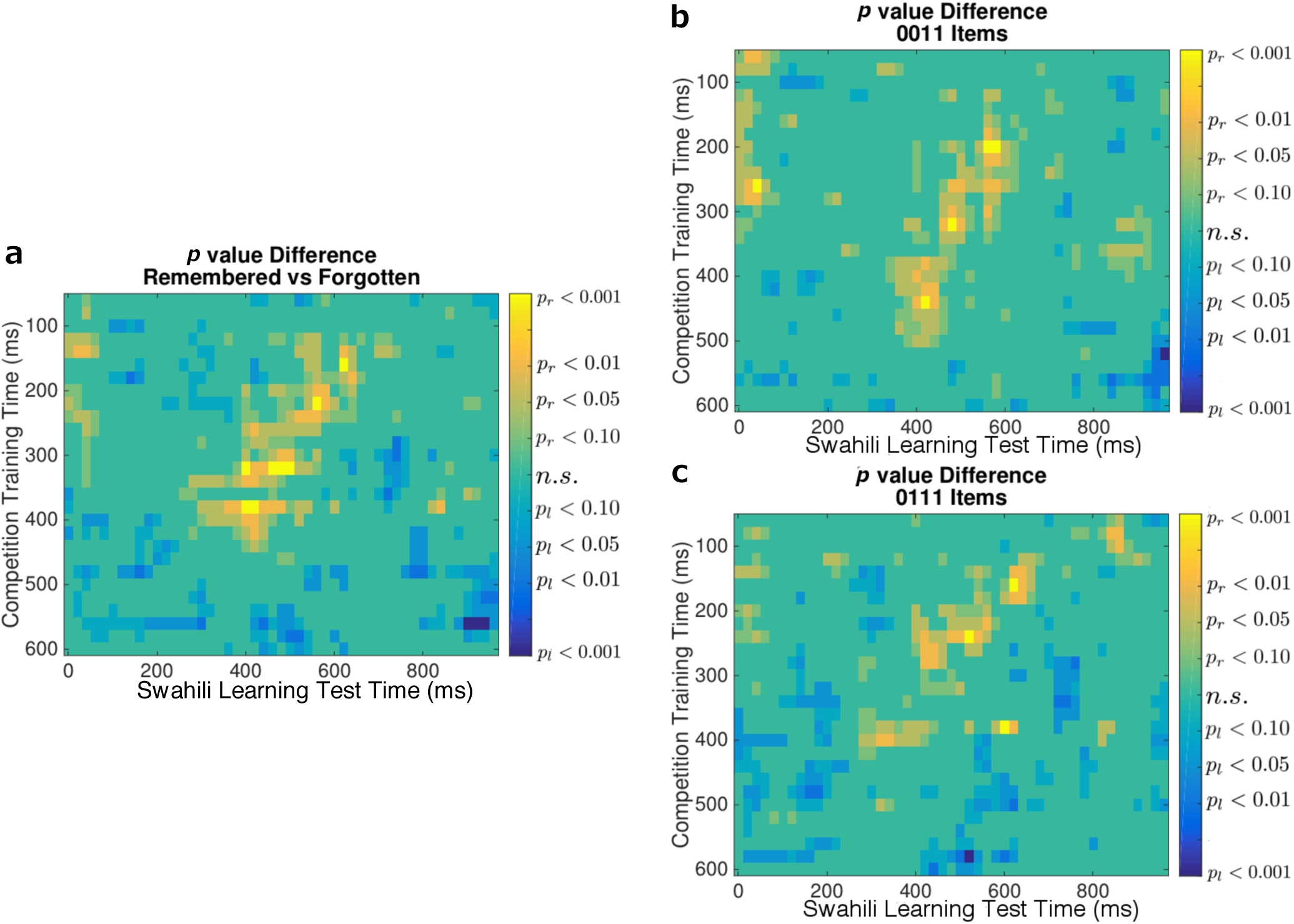
Competition drop analysis with behavior held constant. Comparison of competition drop analysis using all items to that split by behavior during Swahili Learning Task. To evaluate whether the competition drop metric from the EEG data provided addition predictive power over behavior, we re-ran the analysis from Fig. 7 on two distinct subsets of items: items for which subjects were only correct on the last two rounds (0011 items) and items for which subjects were correct on all but the first round (0111 items). **a. One-sided t test results on all trials.** The one-sided t-test result of the difference in competition drop between remembered and forgotten items. Time points for which the drop was larger for remembered items, i.e., where a right-tailed test (*p_r_*) achieved significance, are yellow, whereas time points for which the drop was larger for forgotten items (*p_l_* significant) are blue. **b. Result in subfigure a, computed for 0011 items.** Cluster permutation test significance was *p* = 0.01 for 0011 items. **c. Result in subfigure a, computed for 0111 items.** Cluster permutation test significance was *p* = 0.17 for 0111 items.

Given that our main analysis pooled trials across subjects, both within-subject and across-subject variance could have contributed to the observed relationship between competition drop in Session 2 and subsequent recall in Session 3. That is, it could be the case that subjects who had higher competition drop scores on average had higher levels of Session 3 recall on average; or it could be the case that, within individual subjects, items that had higher competition drop scores were recalled better in Session 3. To address this latter point, we z-scored the competition drop value within subject. This eliminated the influence of between-subject variance. The recomputed grids for all trials, as well as the 0011 and 0111 items (considered separately) are shown in Supplementary Figure S3. Qualitatively, these results strongly resemble the original results, but the predictive power is lessened by z-scoring within subject. This suggests that both across-subject variance and within-subject contributed to our results.

**Figure S3.**
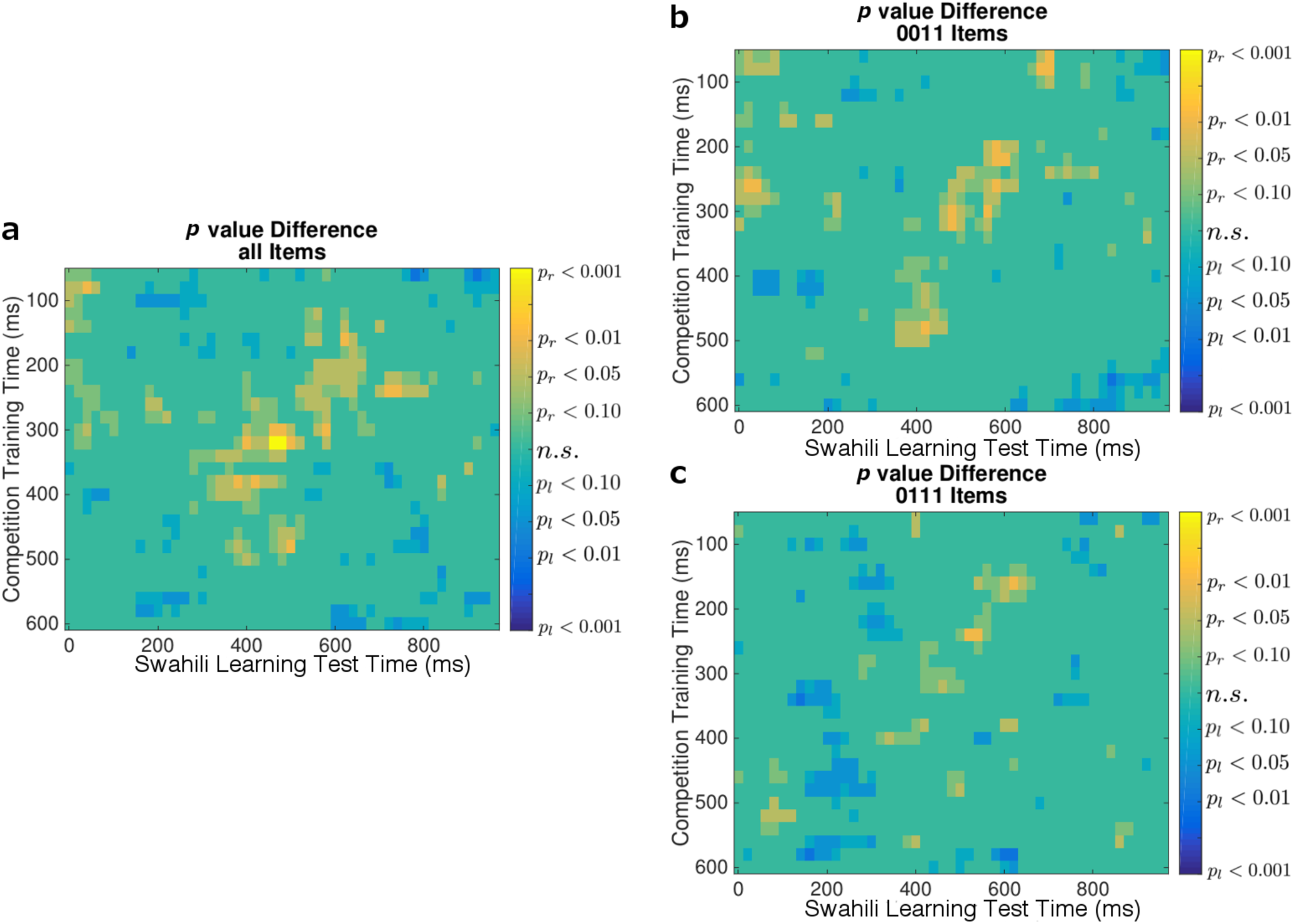
Competition drop analysis with behavior held constant and cross-subject variance mitigated. Results shown in Fig. S2 with within-subject competition drop scores z-scored. **a. One-sided t test results after z-scoring within subject.** Time points for which the drop was larger for remembered items, i.e., where a right-tailed test (*p*_*r*_) achieved significance, are yellow, whereas time points for which the drop was larger for forgotten items (*p*_*l*_ significant) are blue. Note the similarity to Fig. 7b. **b. Result in subfigure a, computed for 0011 items. c. Result in subfigure a, computed for 0111 items.**

Finally, prior work has shown that theta band power in the first 500ms post stimulus onset can predict retrieval-induced forgetting^22^. The results described above were computed using EEG voltages as the input to the classifier; however, the same process can be applied to any set of features derived from the EEG data. To test the ability of theta band power to predict the benefit of retrieval practice, we repeated the above analyses using theta band power values (computed via the Hilbert transform) as the features in our analysis pipeline. We found that high- and low-competition trials in Session 1 could be distinguished with theta band power, shown in Supplementary Figure S4. While there were several timepoints that achieved significance, classifier accuracy was numerically lower than when we used voltage features (see Fig. 5). For subsequent analysis, to facilitate comparison between voltage and theta results, we used the same timepoints for our theta analyses that we used in our voltage analyses (note that this is a superset of the discriminative theta band timepoints). Using these timepoints and testing on the Session 2 data, we did not recover a cluster of timepoints that significantly predicted Session 3 behavior; the results qualitatively resemble the results for voltage features but were not as robust.

**Figure S4.**
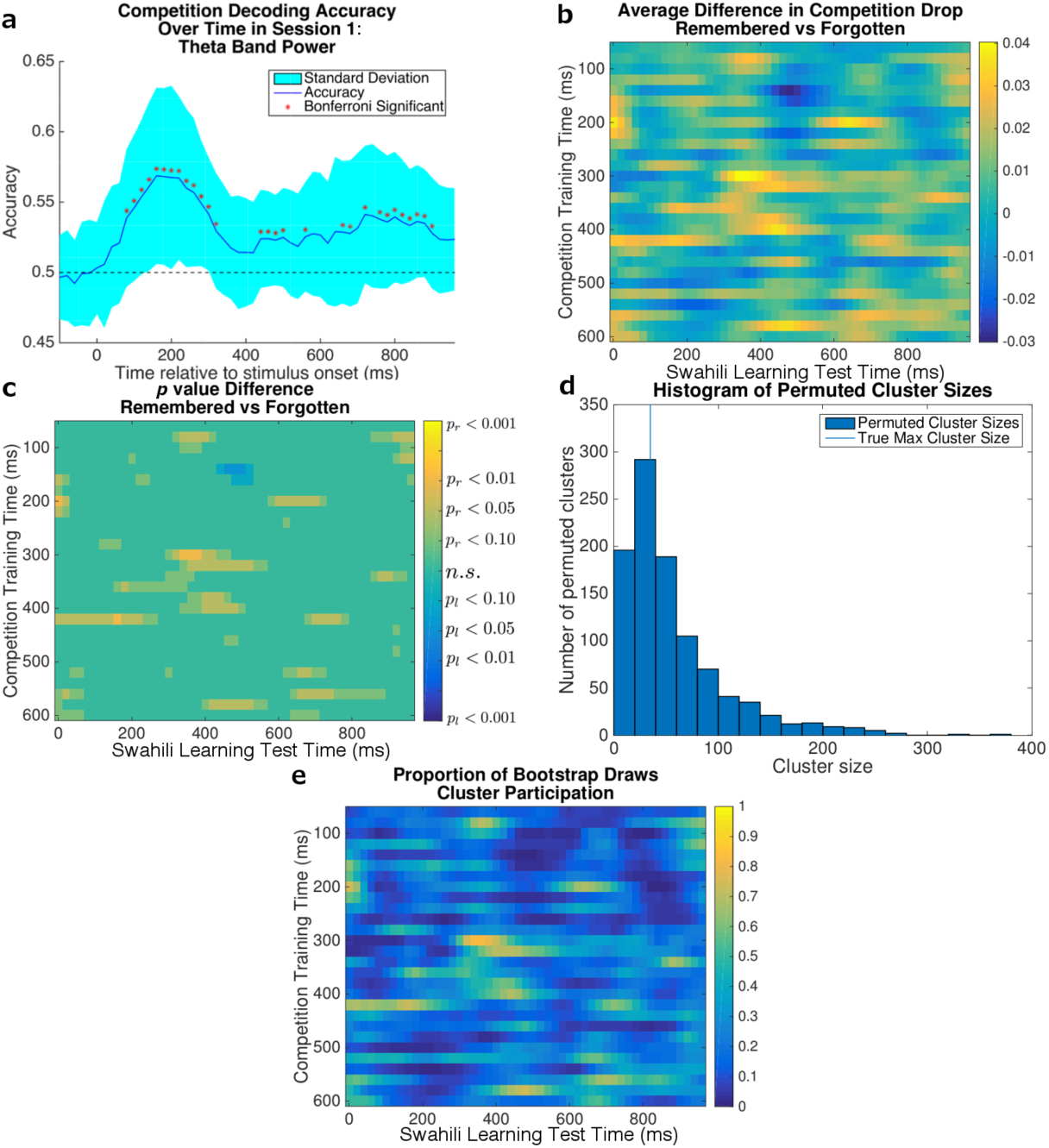
Analyses relating competition drop to subsequent memory, using theta-band power as features. The main analysis summarized in Fig. 5 and Fig. 7, conducted using theta-band power as opposed to voltages. **a. Competition decoding accuracy over time in Session 1** 5-fold cross-validation accuracy at each time point for distinguishing high- and low-competition retrieval states, averaged across subjects. The shaded region gives the standard deviation across the subject means. Chance is 50%. Stars indicate time points that were significant after per-time point permutation test and Bonferroni correction at 0.001 significance threshold. **b. Average difference in competition drop between remembered and forgotten items.** Yellow indicates that remembered items exhibited a larger competition drop than forgotten items. Blue indicates the opposite. **c. One-tailed t-test of the difference in competition drop between remembered and forgotten items.** Grid squares for which the drop was larger for remembered items, i.e., where a right-tailed test (*p_r_*) achieved significance at *p_r_* < 0.10, are yellow; grid squares for which the drop was larger for forgotten items, i.e., where a left-tailed test (*p_l_*) achieved significance at *p_l_ <* 0.10, are blue. The right-tailed test corresponds to our hypothesis that a drop in competition is predictive of memory. No clusters survived multiple corrections. **d. Cluster permutation test histogram** The vertical line indicates the surviving cluster size achieved using the true labels. Each draw in the histogram represents the largest cluster observed in a given permutation of the data. **e. Subject bootstrap results.** Subjects were sampled with replacement; on each draw the cluster analysis was re-run. The color scale on this plot represents the proportion of bootstrap draws in which a point belonged to a significant cluster. A high frequency (yellow) indicates the the point was highly consistent across subjects.

## Discussion

In this study, we trained an EEG classifier to detect neural correlates of competitive retrieval (using a separate *competition localizer* phase), and we used this classifier to track competition across repeated retrieval practice attempts for Swahili-English paired associates. As predicted, we found that the decrease in competition across retrieval practice trials (during learning) was predictive of subsequent recall success one week later.We did not have a strong *a priori* hypothesis about which time windows (post-stimulus onset) would yield a competition readout that predicts subsequent memory, so we swept across a grid of possible time points and found a large cluster of time points (significant after multiple-comparisons correction) that yielded the predicted relationship whereby competition decrease predicts better memory; we did not find any reliable effects in the opposite direction.

Importantly, and unsurprisingly, it was also possible to use participants’ behavior during learning (i.e., the number of times an item was retrieved correctly during retrieval practice trials) to predict memory on the final test one week later. To assess whether our EEG measure of competition drop provided incremental predictive power, beyond what one would get from behavior alone, we ran follow-up analyses where we matched behavioral performance during learning and demonstrated a very similar pattern of results, although significance values were lower (reflecting the decreased number of trials included in these behavior-matched analyses).

Overall, the results are consistent with our hypothesis that benefits of testing on long-term retention are due (at least in part) to reductions in competition^8^. These reductions in competition could arise from multiple mechanisms, e.g., weakening or inhibition of competing memories^13, 25–27^, differentiation of competing memories from the target memory^19, 28–31^, or even integration of competing memories^31–33^. Reduced competition might also owe, in part, to an updated contextual representation stemming from retrieval-based context reinstatement; this, in turn, could have the effect of restricting the search set and allowing for relatively unfettered access to the target material^34,35^. The results of this study do not arbitrate between these possibilities. Furthermore, as noted in the Introduction, researchers have set forth a wide range of explanations for the testing effect other than competition-reduction^1,2^. For example, searching for the correct response on a test leads to the elaboration of semantic associations through which the target material can later be accessed^7,8^. The important point here is that these alternative explanations are not mutually exclusive with the competition-reduction account. Our study was designed specifically to evaluate the competition account and not the other theories; hence, additional work is needed to assess how these different accounts fit together.

Another limitation of the present study is that we did not include a condition where participants simply restudied items instead of doing retrieval practice. Based on prior work with this Swahili-English word-pair-learning paradigm^3^, we expect that this condition would have yielded lower levels of competition (as measured by the classifier) and worse recall compared to the retrieval practice condition. Given that our goal was explaining variance *within* the retrieval practice condition, we decided to omit the restudy condition in order to maximize the number of retrieval practice trials (thereby maximizing our power for detecting within-condition effects).

Our results resemble those of Kuhl et al.^20^, who found (in a between-subjects fashion) that a decrease in a neural measure of competition (in their case: anterior cingulate activity) predicted memory performance. However, there is an important difference between our results and those of Kuhl et al. – while we found a relationship between competition-reduction during retrieval practice and subsequent memory for the practiced items, Kuhl et al. found that decreased anterior cingulate activity was specifically correlated with competitor forgetting (RIF) but *not* improved memory retention for the practiced items^20^. One potential explanation for this discrepancy is that Kuhl et al. used a much shorter retrieval delay than we did: Their experiment included the retrieval practice and final test phases as part of the same fMRI session, whereas we included a 1-week delay between retrieval practice and the final test. Other studies have found that the beneficial effects of testing on target memory are most evident after long delays of the sort that we used here^1,3^. Several mechanisms might be responsible for these delay effects: For example, offline consolidation during the 1-week delay might serve to amplify representational changes (differentiation or integration effects) that are set in motion during retrieval practice^36^; alternatively, competition-reduction might be more important for successful recall when the target is weak (after a long delay) then when it is strong. Regardless of the mechanism, if adding a 1-week delay amplifies the memory benefits of retrieval practice, this might also make it easier to see a relationship between these benefits and competition-reduction during learning.

Another relevant question is how our results relate to those of Hanslmayr et al.^22^, which we used as the basis for our “competition localizer” analyses. One difference between the studies is that our main analysis path (chosen *a priori*) used voltages instead of the theta-power features used by the Hanslmayr et al. study. After completing the analyses with voltage features, we re-ran the main analyses with theta-power features. The results of these analyses, shown in Supplementary Figure S4, conceptually replicate the prior finding^22^ that theta-power features discriminate between high-competition and low-competition items. Competition drop computed using theta-power features during Session 2 did not significantly predict subsequent memory; the pattern of results was qualitatively similar to what was obtained for voltages, but less robust.

In summary: We demonstrated for the first time that decreases in competition (measured neurally) across successive retrieval practice trials are predictive of subsequent retention. This extends prior work by Kuhl and colleagues showing that decreases in competition across trials predict competitor forgetting^20^, by showing that decreases in competition also predict long-term target retention. From a theoretical perspective, this work supports the hypothesis that competition decreases are (at least in part) responsible for the memory benefits of testing. Practically speaking, this result points to potential brain-computer-interface applications: By tracking the decrease in measured competition over time, we may be able to provide online feedback to learners about when a memory as been properly learned. More concretely, learners could be instructed to keep studying a pair until competition levels “bottom out,” indicating that competitors have been suitably neutralized. At present, the EEG signal is still too noisy to provide useful feedback to learners on a trial-by-trial basis, but future improvements in signal-to-noise may make this possible.

## Methods

### Participants

We collected data from 49 participants in total (32 female). Of these, 41 completed all three experimental sessions, while 8 completed only Session 1. Forty (25 female) of these individuals contributed sufficient Session 2 trials (after artifact removal) for subsequent analysis.

Participants were all right-handed native English speakers with normal or corrected-to-normal vision who reported no prior knowledge of Swahili. This study was approved by the Princeton University Institutional Review Board. All participants provided written, informed consent and were monetarily compensated for their time, in accordance with the Princeton University Institutional Review Board compensation guidelines.

### Stimuli

#### Session 1: Competition Localizer Task

Session 1 utilized 19 categories, each of which was linked to eight exemplars of that category (both the category and exemplars were in English). These materials were largely based on the category-exemplar sets used by Anderson, Bjork and Bjork^37^. Exemplars within each category were selected to begin with a unique two-letter string, allowing for selective cueing. The full stimulus set can be found in Supplementary Table S1.

**Table S1.**
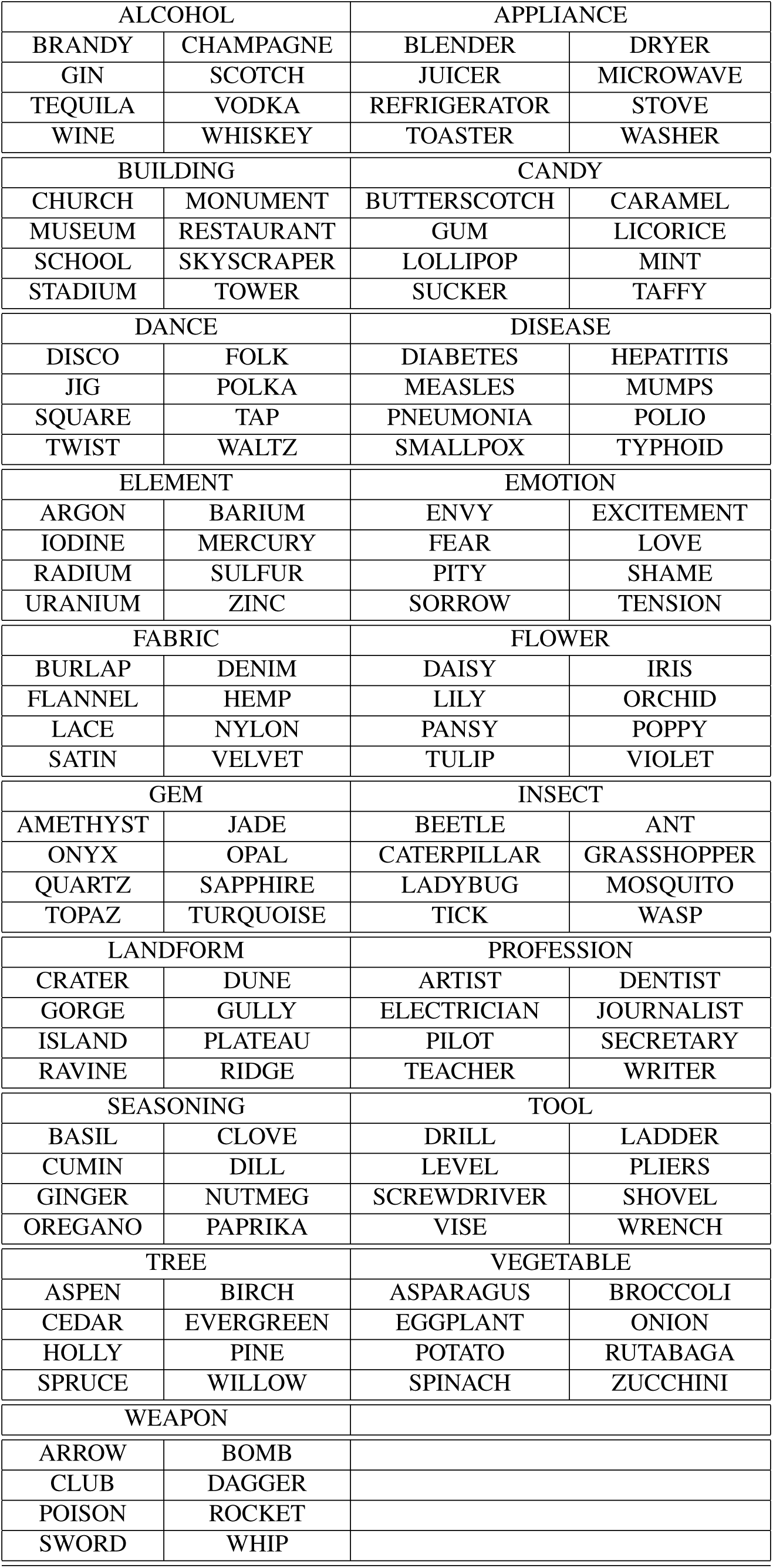
Stimuli for the Competition Localizer Experiment.

#### Session 2: Swahili Learning Task

For Session 2, we used the 40 Swahili-English word pairs from Karpicke and Roediger^3^, augmented with 20 similar Swahili-English word pairs. Pilot studies revealed that participants’ recall performance was nearly perfect when we used only 40 word pairs, leading to a lack of examples of forgotten items for subsequent analysis, hence the increase in number of items from 40 to 60. Stimuli can be found in Supplementary Table S2.

**Table S2.** Stimuli for the Swahili Learning Task.

| Swahili | English | Swahili | English |
| --- | --- | --- | --- |
| BUU | MAGGOT | THELUJI | SNOW |
| FARASI | HORSE | MAITI | CORPSE |
| PUNDA | DONKEY | NYANYA | TOMATO |
| NDOO | BUCKET | MASHUA | BOAT |
| GOTI | KNEE | KAPUTULA | SHORTS |
| ZULIA | CARPET | FUNUNU | RUMOR |
| ELIMU | SCIENCE | BUSTANI | GARDEN |
| LESO | SCARF | DAFINA | TREASURE |
| ADHAMA | HONOR | TUMBILI | MONKEY |
| GODORO | MATTRESS | VUKE | STEAM |
| FAGIO | BROOM | SUMU | POISON |
| CHAKULA | FOOD | KABURI | GRAVE |
| PIPA | BARREL | TABIBU | DOCTOR |
| YAI | EGG | HARIRI | SILK |
| KAA | CRAB | MALKIA | QUEEN |
| POMBE | BEER | ADUI | ENEMY |
| PAZIA | CURTAIN | REMBO | ORNAMENT |
| ZIWA | LAKE | SALA | PRAYER |
| ROHO | SOUL | USINGIZI | SLEEP |
| EMBE | MANGO | ZEITUNI | OLIVES |
| WALI | RICE | MBWA | DOG |
| GARI | CAR | MLIMA | MOUNTAIN |
| SURA | FACE | SAMAKI | FISH |
| MLANGO | DOOR | KOFIA | HAT |
| MENDE | BEETLE | UPEPO | WIND |
| KOLEO | SHOVEL | SHAKA | PROBLEM |
| MTO | PILLOW | JESHI | ARMY |
| NJIA | ROAD | MAJI | WATER |
| NYOKA | SNAKE | RINDA | DRESS |
| UMBU | SIBLING | MAZIWA | MILK |

#### Session 3: Swahili Follow-up Test

In Session 3, participants were tested on the Swahili words presented in Session 2 using Google Forms.

### EEG Recording

A Biosemi ActiveTwo EEG Data collection system was used, in a shielded room featuring an RF/EMI Faraday cage built by Universal Shielding Company. The acquisition software was ActiView 7.07 on a 64-bit Windows workstation. A 64-channel configuration and a sampling rate of 512 Hz was used, along with 6 flathead electrodes (2 to record eye blinks, 2 for bilateral eye movement, and 2 on each mastoid for referencing).

EEG data were preprocessed using EEGLab^38^. First, the data were visually inspected for major artifacts and those sections of data removed. Then the data were band passed between 2 and 200Hz, with a notch filter applied at 60Hz to remove line noise. The electrodes were re-referenced using the whole scalp, then the data were parsed into epochs using stimuli onset and the epochs were baseline corrected. To remove eye-blink artifacts, the top two independent components were removed, after being visually inspected. After preprocessing, 2170 trials out of a total of 2400 (pooled across participants) from Session 2 survived for subsequent analysis. On average 295 out of 304 trials (per subject) from Session 1 survived.

For the experiment using theta band power, theta band was defined as between 4 and 8Hz, inclusive, and was extracted using a Hilbert transform after the electrode referencing step. The data were then parsed into epochs without baseline correction or blink removal.

### Experimental Design

There were three experimental sessions: see Fig 1 for a schematic. In Session 1, participants were shown the high/low competition retrieval stimuli while EEG was recorded. In Session 2, participants were given four study and test rounds for 60 novel Swahili words while EEG was recorded. Session 3 consisted of a cued-recall test of the Swahili words studied in Session 2, followed by a questionnaire. No EEG recording took place in Session 3.

#### Session 1: Competition Localizer Task

Session 1 was composed of two parts: an initial study phase and then a test phase. In the study phase, the participant was shown all the category-exemplar pairs in the experiment in one block. Each pair was presented for 2s; a fixation cross was presented for 1s between each pair. The category was presented above fixation and the exemplar was presented below fixation. Participants were instructed to try to remember the category-exemplar pairs. The test phase of the experiment consisted of four blocks in which the participant was presented with part of a category-exemplar pair. In each trial, the participant was presented with either the category or the exemplar. The other component of the pair was only partially presented, with the first two letters followed by two dashes (not diagnostic of the length of the word). Each partial pair was presented for 3s, with 1s of fixation between trials. Participants were instructed to silently fill in the blank with the correct word. Trials in which the category was provided required retrieval of the exemplar from many similar competitors, and are designated “high-competition trials.” Trials in which the exemplar was provided were assumed to reflect a relatively low level of retrieval competition and are referred to as “low-competition trials.” Participants were given the opportunity to rest between each block, as well as after the study block. Each block was composed of equal numbers of high-competition and low-competition trials, randomly interleaved. All 19 categories *×* 8 items *×* 2 conditions possible partial pairs were each shown once in the test phase.

#### Session 2: Swahili Learning Task

If time permitted, Session 2 was run immediately following Session 1; if not, participants returned within one week to complete Session 2. This session consisted of four rounds, with each round consisting of a study block and test block. In the study blocks, the participant read the Swahili-English word pairs in a random order. Each pair was presented for 2s, and a fixation cross was presented for 1s between trials. In the test blocks, participants were shown just the Swahili words (in a re-randomized order) and asked to respond with the English translation. Participants were asked to think of the answer silently for 2s prior to typing it; a delay was programmed such that keystrokes occurring within 2s of the Swahili word presentation would not appear. When finished, the participants pressed “enter” to move to the next Swahili word, and were free to skip words if need be. They were instructed not to spend too much time worrying about the response and to skip the question if unsure. Participants were given the opportunity to rest between each block. Subjects were asked not to review Swahili vocabulary between Session 2 and Session 3.

#### Session 3: Swahili Follow-up Test

One week after Session 2, participants returned to the same room to take a final test on the Swahili words presented in the previous session. Participants took the test in the same chair and with the same computer setup as the previous sessions, but they were not outfitted with an EEG cap as no electrophysiological data were recorded during this session.

### Behavioral Data Analysis

For each Swahili-English word pair, each participant produced 4 behavioral datapoints in Session 2 (correct or incorrect recall, for each test round). We examined whether the response pattern of each individual item over the four rounds of Session 2 could predict which items were remembered or forgotten in Session 3. To evaluate the predictive power of the Session 2 responses, we used *L*_2_ penalized logistic regression. Each item was represented by a four dimensional binary vector indicating whether the participant answered correctly in each of the Session 2 rounds. These were the features used to predict the label, a binary scalar indicating whether the item was remembered in Session 3. To establish significance, a permutation test was conducted with 1000 permutations.

### EEG Predictive Modeling Framework

Our modeling framework can be outlined as follows: First, we located the times relative to stimulus onset in Session 1 at which high competition and low competition neural states were significantly distinguishable by a logistic regression classifier, with a suitably large effect size (accuracy *>* 1 SD above chance). Then we trained our classifier on the data from each of those time points separately and applied it to each time point post-stimulus-onset, for each retrieval trial in Session 2. This created, for each of the four testing rounds for a given item, a grid of competition estimates (where the grid is defined by the time point where the classifier was trained in Session 1, and the time point where the classifier was applied in Session 2). We computed a *competition drop* score for each item by subtracting the competition estimate for the last round from the competition estimate for the first round in which the participant answered correctly; this was done separately for every grid square. We then used a t-test to establish, for each grid square, whether competition drop scores in that grid square were predictive of subsequent memory, and corrected for multiple comparisons. Each of these steps is described in more detail below. This framework was applied both to voltage data and to theta band power data.

### Classifying High Competition and Low Competition Neural States

For each participant, we had Session 1 EEG data corresponding to roughly 300 trials (depending on visual artifact removal), with half of these trials consisting of high competition retrieval and half consisting of low competition retrieval. The EEG data consisted of a multidimensional timeseries of voltage (or theta-band power) at each channel and each timepoint. For each time point, we averaged the points for the next 50ms, creating a 64-dimensional feature vector representing the average activity in each channel for the 50ms period starting at that time point. This feature vector was input into a binary *L*_2_ penalized logistic regression classifier. The data were divided into five cross-validation folds. For each fold and time point, the classifier was trained on four of the folds and tested on the fifth. Thus we generated time series of classification accuracy for decoding high vs low competition states from the EEG data. This process was applied separately for each participant, and the accuracy timeseries was averaged across participants.

Significance and population-level reliability were measured using a within-participants permutation test with 100 permutations, and the within-participants *p* values were combined by treating participant as a random effect. Each participant’s *p* value was converted into a *z* value via the inverse normal transform. A group *p* value was generated by a one sample *t*-test between these *z* values and zero. The group *p* value at each timepoint was corrected for multiple comparisons using Bonferroni correction for an initial group *p* value of 0.001 divided by the total number of timepoints (55).

To generate the representational dissimilarity matrix over the weight vectors, we computed the cosine distance of the weight vectors for each pair of timepoints. To examine the neural representation underlying a given weight vector, we used the method described by Haufe et. al^24^, and averaged the resulting patterns over subjects and over the timepoints of interest.

### Tracking Competition Level during Cued Recall

For each participant, a logistic regression classifier was trained to distinguish high-competition from low-competition trials using all trials from Session 1; a separate classifier was trained for each timepoint that was significant (and where accuracy was greater than 1 SD above chance) in the Session 1 group analysis. Each participant’s Session 2 data contained four retrieval rounds for each of the 60 words. These data, like the Session 1 data, form a multidimensional timeseries, and we again looked at each 50ms window average separately, creating a 64-dimensional feature vector to which the trained classifier could be applied. Logistic regression output, when not thresholded into classes, can be treated as the estimated probability of a sample belonging to the “1” class. We refer to this output as the “competition probability,” the probability that high competition retrieval is occurring. For each of the four retrieval rounds and each of the 60 items, our procedure generated a grid of this competition probability, with each point in the grid consisting of a training time from Session 1 and a test time from Session 2.

### Predicting Successful Learning

The main goal of our analysis was to assess whether the decrease in competition probability across Session 2 rounds predicted subsequent recall in Session 3. To achieve this goal, we first computed (for each item, and for each grid square) a competition drop score: the competition probability for the first round in which the participant responded correctly minus the competition probability for the last round (round 4). If the participant never responded correctly or did not respond correctly in round 4, the item was dropped from further analysis. This reduced the total number of trials from 2170 to 1550. Next, for each grid square, we took the competition drop scores for all trials (pooled across all subjects) where the item was subsequently remembered, and compared them to the competition drop scores for all trials where the item was subsequently forgotten. The distributions of competition drop scores for subsequently-remembered and subsequently-forgotten items were Gaussian according to the Kolmogorov-Smirnov test^39^. Thus, we could use a one-tailed *t*-test to measure how well the competition drop distinguishes remembered and forgotten items. This generated a grid of *p* values for every grid square. Note that, unto themselves, the *p* values for this test tend to be liberal (insofar as it is a fixed-effects test), but this is remedied by the cluster-correction procedure described below.

To correct for multiple comparisons, we used a variant of the cluster permutation test to compute family-wise error^40,41^. First, we thresholded the *p* values at 0.10. We then computed the sizes of all clusters in the grid, where a cluster was defined as a set of contiguous grid squares that all had *p* values less than 0.10, and the cluster size was the number of grid squares belonging to the cluster. Contiguous was defined as immediately adjacent timepoints, not counting diagonal adjacency.

We then permuted the assignment of items to the subsequently-remembered and subsequently-forgotten clusters, recomputed the grid of *p* values, and thresholded again. For each permutation, the size of the maximum resulting cluster was stored. This was repeated 1000 times. The true clusters were then compared to the histogram of permuted cluster sizes and a family-wise corrected *p* value was generated. Only clusters with a family-wise *p* value less than 0.05 were considered significant.

To measure the population-level reliability of the surviving cluster, we sampled participants with replacement and recomputed the test statistic over all the timepoints. We then looked at the fraction of bootstrap draws in which each timepoint was part of a significant cluster. If the effect is present throughout the population, this should be close to 1 for all points in the true **cluster. If the effect is instead dominated by only a few participants, the fraction would be much lower.We computed 1000** bootstrap draws.

## Data Availability Statement

The datasets generated during and analyzed during the current study are available via the Princeton University dataspace: http://arks.princeton.edu/ark:/88435/dsp015138jh55f. Code used for data collection and analysis are available at https://github.com/PrincetonCompMemLab/comp-eeg. Code used to generate Figures 7 and Supplementary Figures S2, S3, and S4 is available for manipulation via CodeOcean: https://codeocean.com/2018/03/30/competition-eeg-v3/

## Acknowledgements

This work was supported by the National Science Foundation Graduate Research Fellowship under Grant No. DGE1252522, and National Institutes of Health grants No. R01HD075328 and No. R01MH069456.

## Author contributions statement

N.S.R., J.C.H., and K.A.N. conceived the experiment. N.S.R. and P.P. collected the data. N.S.R. analyzed the results. N.S.R., K.A.N., and J.C.H. wrote the manuscript. All authors reviewed the manuscript.

